# A VTA-pontine GABA pathway biases locomotor direction via local and distal inhibition

**DOI:** 10.64898/2026.02.17.706301

**Authors:** Cristian González-Cabrera, Rukhshona Kayumova, Ezia Guatteo, Nicola Berretta, Nicola B. Mercuri, Trinidad Montero, Miquel Vila, Pablo Henny, Matthias Prigge

## Abstract

Locomotor direction in mammals is implemented by descending circuits, yet how midbrain selection systems bias directional motor output remains unclear. Here we define a projection-defined inhibitory pathway from the ventral tegmental area to the oral pontine reticular nucleus (VTA_PnO_) whose activation is sufficient to drive backward locomotion. These TH*^−^* VTA neurons form monosynaptic GABA_A_ synapses locally while projecting to PnO, establishing a dual local–projection inhibitory architecture. Somatic activation reliably induced backward locomotion, and selective stimulation of VTA_PnO_ terminals reproduced the effect. Pathway recruitment produced a rapid transient increase in dopaminergic single-unit activity and frequency-dependent increases in dopaminergic population calcium signals in awake mice. During forced locomotion, chronically recorded VTA_PnO_ neurons were preferentially engaged during reverse compared to forward rotations. Together, these findings reveal a projection-defined midbrain pathway that biases locomotor direction through coordinated local inhibition and distal engagement of a brainstem premotor node.

## Introduction

The ventral tegmental area (VTA) is a midbrain hub for motivational and behavioral-state selection and reinforcement, with pronounced cellular and circuit heterogeneity. While dopaminergic (DA) neurons anchor models of reward prediction, addiction, and valence [1–4], GABA neurons in the VTA exert strong local control over dopaminergic activity. Inhibitory signaling regulates DA excitability and can shape firing patterns, including via local disinhibitory circuits [5,6]. Consistent with this local control, optogenetic manipulation of VTA GABA neurons is sufficient to bias behavior, including aversive learning and reward consumption [7,8]. Beyond these local computations, it is well established that the VTA contains long-range GABAergic and glutamatergic projection neurons that innervate diverse downstream targets [9–12]. However, recent work challenges a strict local-versus-projection dichotomy by showing that long-range VTA projection neurons can also form intra-VTA synapses, revealing a dual local-and-projection architecture with parallel influence on both: local VTA circuits and downstream targets [13]. Additionally, molecular and anatomical studies emphasize that VTA output channels are functionally specialized and projection-defined [14,15]. Here, we test the functional consequences of such dual local-and-projection architecture for the control of locomotor direction. Locomotor direction in mammals is implemented by descending premotor circuits in the pontomedullary reticular formation; within this axis, the oral pontine reticular nucleus (PnO) projects to the spinal cord and interfaces with reticulospinal controllers [16–18]. Classic pharmacology demonstrated that broad elevations of monoaminergic tone can elicit backward walking and circling, and that receptor blockade suppresses these behaviors [19,20]. Together, this places the VTA, an area with modulatory reach and projection-defined outputs, in anatomical and functional proximity to brainstem direction controllers. Yet, it is unknown if a distinct VTA cell class can selectively bias locomotor direction via downstream nodes. Here, we address this gap by testing a defined VTA to PnO pathway. Within this framework, we tested whether a projection-defined VTA pathway with a dual architecture can engage PnO to bias locomotor direction. Using single-cell juxtacellular labeling, we identified TH*^−^* (non-dopaminergic) VTA projection neurons, including VGAT+ (GABAergic) cells with local collaterals and found a subset that targets the PnO. Focusing on this VTA_PnO_ population with a local-and-projection architecture, somatic activation was sufficient to elicit backward locomotion, and PnO-terminal activation was likewise sufficient, showing a timing difference consistent with convergent local and projection-mediated engagement. Recruiting this pathway elevated VTA DA population signals with a brief DA single-unit increase in acute optogenetic experiments, whereas in chronic recordings during forced backward locomotion, opto-tagged VTA_PnO_ neurons showed distinct, temporally structured responses. These results identify a VTA_PnO_ inhibitory pathway that links midbrain selection circuitry to brainstem premotor command via a dual local-and-projection architecture.

## Results

### VTA outputs previewed by bulk labeling and resolved at single-cell resolution

To examine whether VTA GABAergic projection neurons form dual local–projection architectures, we first mapped their long-range outputs. We injected AAV::CAG-DIO-mNeon into the VTA of Gad2-Cre mice. In line with earlier reports that VTA GABAergic neurons send extensive efferents beyond local circuits [9,11,12], we observed widespread axonal labeling across forebrain and brainstem regions, including nucleus accumbens, lateral hypothalamus and PnO (Fig. 1a,b, Supplementary Fig. 1a). To isolate long-range projecting GABAergic neurons from local connecting neurons we combined juxtacellular recordings and Neurobiotin labeling with post hoc neurochemical and target classification (n = 42; Fig. 1c,d, Supplementary Fig. 1b,c; [21]). Notably, all recovered TH*^−^* neurons (32/32) extended axons outside the VTA, supporting their classification as bona fide projection neurons. The majority of them (62%) also formed local collaterals in the VTA, consistent with a dual projection architecture (Fig. 1d,e), [11,13]. Cell type differences were also evident in spike timing variability (CV2) and waveform. Local-and-projection (LP) TH*^−^* neurons fired more irregular than projection-only (PO) TH*^−^* neurons (Supplementary Fig. 1d). Within both classes, firing rate covaried inversely with spike-time variability (Supplementary Fig. 1d). Spike waveforms from TH*^−^* neurons were significantly narrower (shorter duration) than those of identified DA neurons (Fig. 1e) consistent with established in vivo criteria [22,23]. Baseline firing rate of recorded neurons did not show any difference between groups (Fig. 1e). A 15s hindpaw pinch differentially modulated identified VTA neurons: TH+ cells showed predominant decreases in firing rate, whereas TH*^−^* neurons exhibited heterogeneous rate changes (both increases and decreases), yielding a significant population-level contrast (Supplementary Fig. 1e). Burst dynamics likewise diverged, TH*^−^* units tended to increase or maintain bursting, while TH+ neurons were largely unchanged or reduced (Supplementary Fig. 1f). Because juxtacellular fills are passive and not always enough to yield extensive axonal labeling, identified targets represent lower-bound estimates [21]. Yet the coexistence of local and distal (projection) axons observed in single-labeled neurons demonstrates that TH*^−^* VTA-projection neurons can form both local and distal axons, consistent with a dual local-projection architecture [11,13]. Among the LP neurons, a subset projected to the PnO (12%, 4/32 TH*^−^*), identifying a VTA_PnO_ subpopulation (Fig. 1b), which we next targeted ex vivo to test dual architecture motifs.

**Figure 1:**
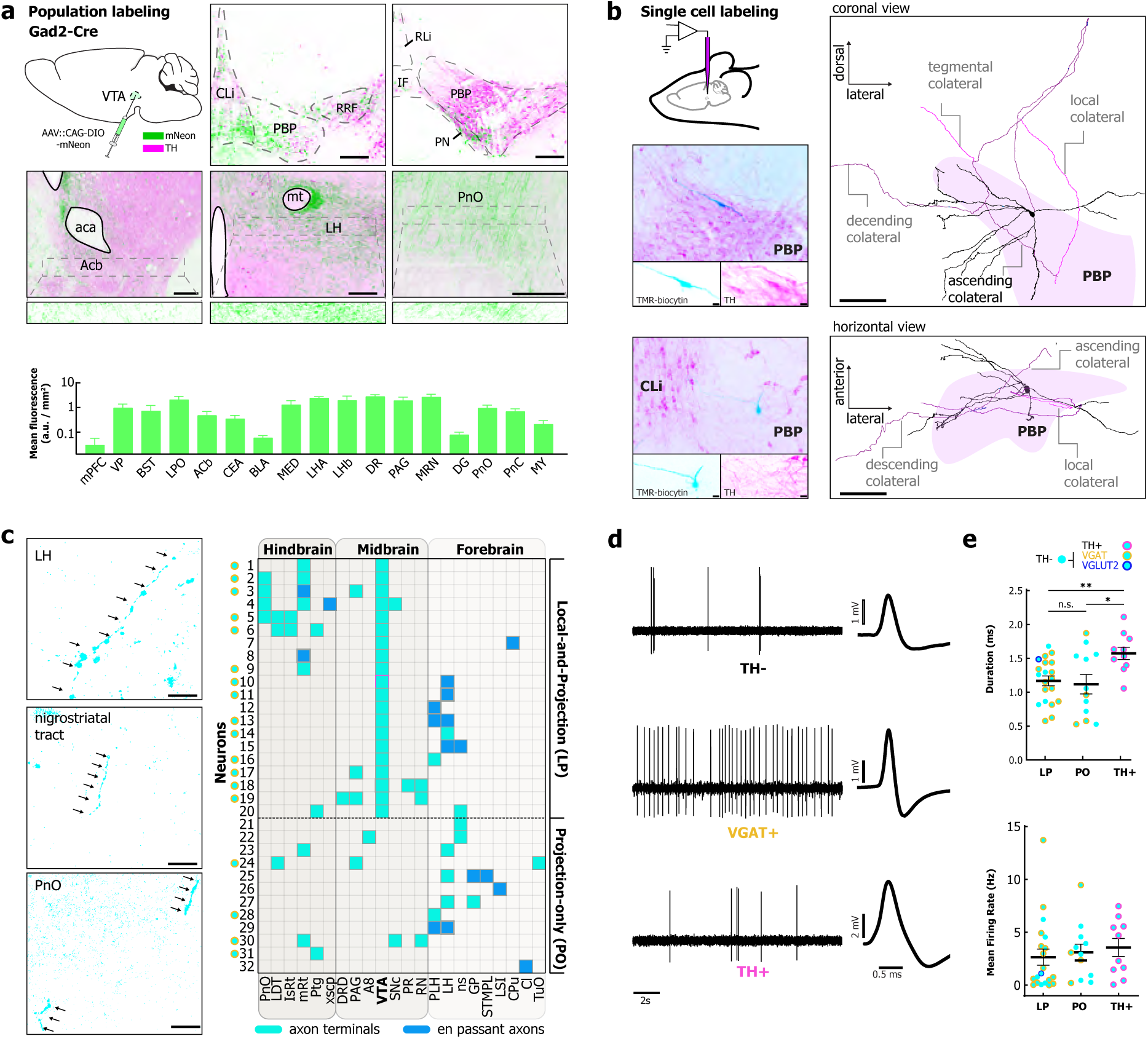
TH*^−^* VTA neurons combine local collaterals with long-range outputs. **a,** Population labeling and projection overview. Upper panel, population strategy (example shown in Gad2-Cre): a single VTA injection of AAVDJ-CAG-DIO-mNeon labels GABAergic VTA neurons. Representative coronal sections illustrate reporter expression (green) in VTA subfields (CLi, PBP, RRF, IF, RLi, PN) and long-range labeling in nucleus accumbens (Acb), lateral hypothalamus (LH), and the oral pontine reticular nucleus (PnO). Magenta: TH immunostaining. aca: anterior commissure, mt: mamillothalamic tract. Bottom panel, projection strength across selected targets expressed as mean fluorescence. **b,** Single-cell juxtacellular recording and labeling. Left, biocytin-filled neurons with post-hoc TH immunostaining identify TH^+^ and TH*^−^* cells. Right, coronal and horizontal 3D reconstructions from a TH*^−^* neuron reveal local VTA collaterals together with ascending and descending long-range collaterals. **c,** Axonal phenotypes from identified neurons. Left, examples of axon terminals and en passant profiles in target structures (arrows). Right, per-neuron (*n* = 32; recorded TH*^−^* neurons) matrix summarizing targets; cyan = terminals, blue = en passant. Neurons are grouped by whether intra-VTA collaterals were present (Local-and-Projection; LP) or absent (Projection-only; PO). Green circles denote GABAergic cells. **d,** Electrophysiological signatures: representative traces and average waveforms from TH^+^, TH*^−^*, and VGAT^+^ neurons. **e,** Spike duration was shorter in TH*^−^* than in TH^+^ neurons (TH^+^ vs. LP: *p* = 0.0023; TH^+^ vs. PO: *p* = 0.0185; one-way ANOVA, FDR-corrected post hoc tests). Mean firing rate did not differ between cell groups (Kruskal–Wallis test, FDR-corrected post hoc tests). LP: *n* = 20; PO: *n* = 12; TH^+^: *n* = 10. Scale bars: A and C: 200 *µ*m; B: 20 *µ*m and 200 *µ*m.

### A VTA GABA subpopulation combines long-range output with local inhibition

To test whether VTA_PnO_ neurons make local inhibitory synapses within VTA, we expressed ChR2 selectively in PnO-projecting VTA neurons by injecting AAVretro carrying cre-recombinase into PnO and AAV5::hSyn1-DIO-ChR2-mCherry into VTA of C57/BL6 mice (n = 5; Fig. 2a). After more than 4 weeks, we euthanized the animals, prepared acute VTA slices for extracellular single unit and intracellular patch-clamp recordings, and distinguished putative dopaminergic from putative non-dopaminergic neurons by their electrophysiological signatures (AP width, Ih current, and AHP area; Fig. 2b; Supplementary Fig. 2a,b) [23–25]. Brief photostimulation evoked optical IPSCs in putative dopaminergic and GABAergic neurons in the VTA (Fig. 2a). Repetitive stimulation trains at various frequencies, evoked reliable oIPSCs with decreasing amplitude (Supplementary Fig. 2c). These currents were eliminated by GABA_A_ receptor antagonists, confirming inhibitory identity. Synaptic latencies calculated from light-on under aCSF conditions were below 4-ms typical of optical evoked synaptic responses, while addition of TTX and 4-AP did not abolish oIPSCs, but shifted latency to circa 5 ms (Fig. 2c).Together these data show that VTA_PnO_-projecting neurons also provide monosynaptic local GABA_A_ inhibition, demonstrating a functional intra-VTA inhibitory action of these pontine projecting neurons [5–8, 13].

**Figure 2:**
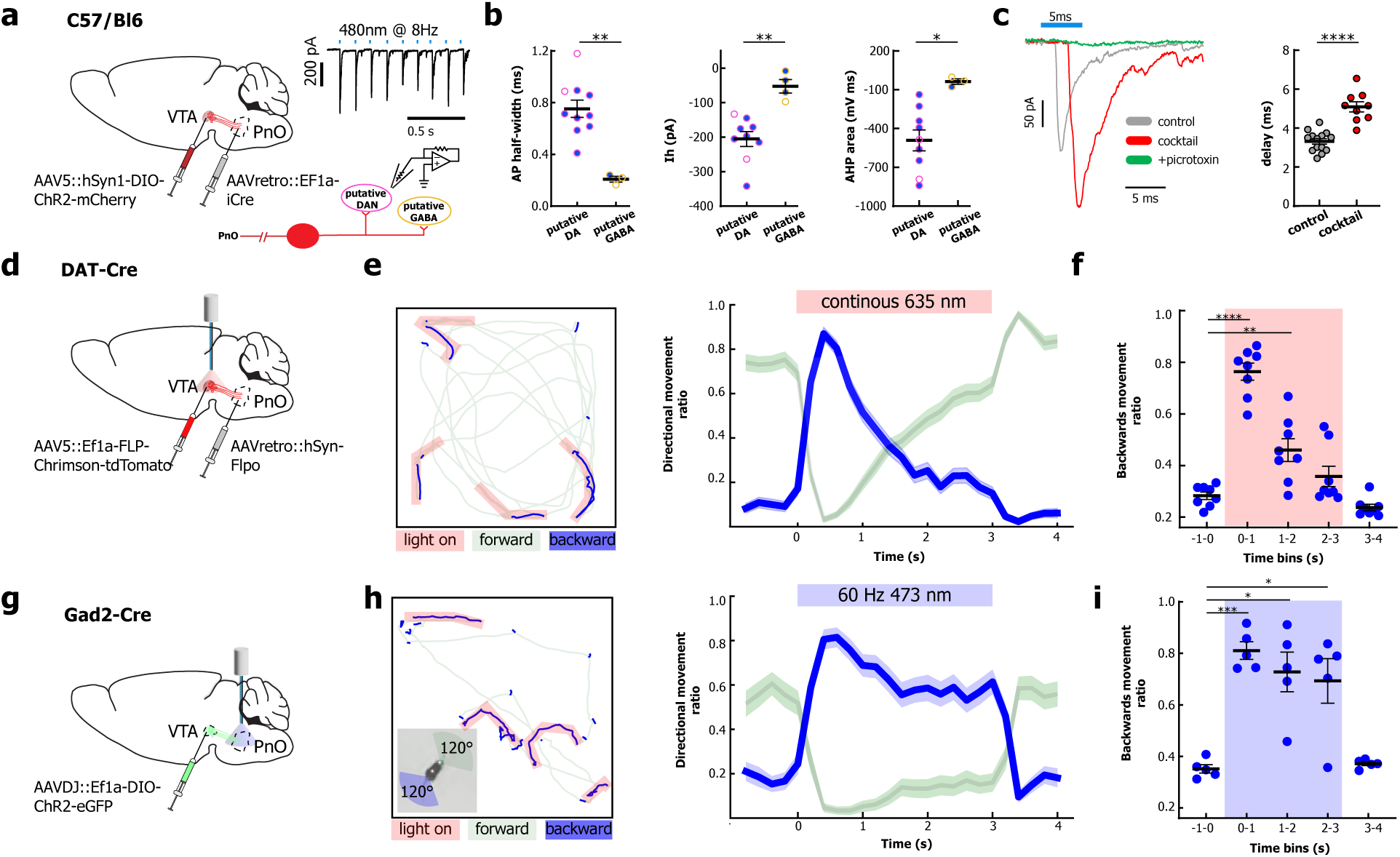
Local VTA effects of VTA_PnO_ neurons and pathway-driven backward locomotion. **a,** Circuit targeting and local recordings. Double-injection strategy to label VTA_PnO_ neurons: retroAAV-Cre in PnO and Cre-dependent ChR2-mCherry in VTA. Representative light-evoked IPSCs (5-ms pulses; 480 nm; example 8 Hz train); schematic shows recordings from unlabeled VTA neurons while optical stimulation activates ChR2-positive VTA_PnO_ local terminals. **b,** Intrinsic properties (AP half-width, *I_h_*, AHP area) separate putative DA from putative GABA neurons (*p* = 0.0012, *p* = 0.0012, *p* = 0.0105, respectively; unpaired *t*-tests). Empty circles indicate neurons non-responsive to light. **c,** Single-pulse IPSCs persist in control and cocktail conditions (CNQX + D-AP5 + 4-AP + TTX) but are abolished by picrotoxin, indicating monosynaptic GABA_A_ transmission from VTA_PnO_ terminals onto local VTA targets. IPSC peak delay is increased in cocktail (5.09 *±* 0.26 ms, *n* = 9) vs. control (3.32 *±* 0.14 ms, *n* = 14; unpaired *t*-test, *p <* 0.0001). **d,** Schematic of VTA_PnO_ somatic activation strategy: retroAAV-Flp in PnO and AAV-FLP-Chrimson-tdTomato in VTA; optic fiber in VTA. **e,** Pathway activation biases locomotor direction in open-field. Left, example open-field track (forward = green, backward = blue; red = 3 s 635-nm continuous light stimulation). Right, population time course of % backward vs. forward movement (*n* = 8). **f,** Backward movement ratio (proportion of backward movement) in 1-s bins (colored area: light stimulation at 1–3 s). Backward ratio exceeded baseline at 0–1 s and 1–2 s (*p <* 0.0001 and *p* = 0.0088, respectively), dropped to baseline at 2–3 s (*p* = 0.086), and fell below baseline at 3–4 s (*p* = 0.04; one-way ANOVA with repeated measures, FDR-corrected post hoc tests). **g,** Schematic of VTA_PnO_ terminals activation strategy: Gad2-Cre: VTA AAV-DIO-ChR2-EGFP in VTA of Gad2-Cre mice; optic fiber in PnO. **h,** Left, same as **e** with red = 3 s, 60 Hz, 470-nm light stimulation. The square insert at the bottom depicts classifications of forward vs. backward movements (for details, see Methods). Right, population time course of % backward vs. forward movement (*n* = 5). **i,** Same as **f** for Gad2 mice (colored area: light stimulation at 1–3 s). Backward ratio was above baseline at 0–1 s, 1–2 s, and 2–3 s (*p* = 0.0006, *p* = 0.0194, and *p* = 0.03, respectively) and not different from baseline at 3–4 s (*p* = 0.33; one-way ANOVA with repeated measures, FDR-corrected post hoc tests).

### Optogenetic activation of the VTA_PnO_ pathway is sufficient to drive backward locomotion

Having identified VTA_PnO_ neurons as a VTA subpopulation exhibiting a dual local-projection architecture, we investigated behavioral output using two complementary strategies. First, we used a projection-specific strategy to express a Flp-dependent red-absorbing channelrhodopsin Chrimson in VTA neurons that project to PnO, then stimulated their somata in VTA via bilateral optic fibers (n = 8; Fig. 2d). Three seconds of continuous illumination with red light in VTA (635nm, 5mW) evoked clear bouts of backward walking in an open-field (Fig. 2e, left; example track: forward in green, backward in blue; light-on in red). The population PSTH (peristimulus time histogram) showed a sharp onset peak in the backward movement ratio (blue) that declined across the 3-s light window and fell below baseline after offset, while forward movement (green) shows inverse relation (Fig. 2e, right). In 1-s time bins, the backward movement ratio exceeded baseline at 0-1 s and 1-2 s and was comparable with the baseline at 2-3 s but below it at 3-4 s (Fig. 2f). Secondly, to isolate the contribution of GABAergic projecting neurons in the PnO from their local collaterals in the VTA, we expressed blue-absorbing ChR2 in the VTA in Gad2-Cre mice (n = 5; Fig. 2g). To prevent presynaptic depolarization block [26,27], we used a 60 Hz blue-light stimulation protocol to drive GABAergic terminals in the PnO (Fig. 2h left, Supplementary Fig. 2d right). This stimulation likewise triggered backward walking and yielded a transient increase in the backward movement ratio that returned to baseline after the stimulus (Fig. 2h right). In 1-s time bins, the backward movement ratio was higher than baseline at 0-1 s, 1-2 s, and 2-3 s but indistinguishable from baseline at 3-4 s (Fig. 2i). Together, both manipulations were sufficient to trigger backward locomotion, but with distinct temporal profiles: somatic VTA stimulation produced a strong onset peak that progressively declined during the 3-s light-on window and dipped below baseline after offset, whereas presynaptic activation of GABAergic VTA_PnO_ terminals produced a constant backward locomotion that persisted throughout the 3-s stimulus window, and returned to baseline thereafter (Supplementary Fig. 2e). Latency to peak backward movement was comparable between manipulations (Supplementary Fig. 2e). By contrast, the time to maximum speed for VTA somatic stimulation was less than for PnO terminals (Supplementary Fig. 2f). Backward movement scaled with stimulation frequency: it was minimal at *≤* 20 Hz and robust at *≥* 60 Hz and during continuous light. A per-trial mixed-effects model on Δ (backward ratio) showed a significant linear trend across frequencies (*n* = 8 mice, 14 trials/mouse). Post-hoc pairwise tests with Holm’s step-down correction indicated that 30–60 Hz elicited greater responses than 20 Hz, whereas 60 Hz vs. continuous light were not significantly different for VTA somatic stimulation (Supplementary Fig. 2d).

### Optogenetic activation of VTA_PnO_ elevates dopaminergic population signals in vivo

To directly quantify how VTA_PnO_ activation shapes local dopaminergic activity in the VTA, we combined our PnO-specific optogenetic labeling strategy with fiber photometry of dopaminergic populations (GCaMP8f in DA neurons) in DAT-Cre mice (n = 5; Fig. 3a). Using a short baseline (-1.0 to -0.1 s) and a peristimulus window (0-3 s), we calculated within-opsin contrasts (Stim vs. Pre) for 20 Hz and continuous stimulation. Based on behavior, continuous stimulation was chosen as the positive regimen (reliably drives backward walking), whereas 20 Hz and below (5-10 Hz) were selected as a boundary, behavior-negative controls (Supplementary Fig. 2d: 30 Hz and above produced backward locomotion). Within-opsin pre-post comparison was significant at continuous stimulation, with 5-20 Hz yielding non-significant differences (Fig. 3b). Similarly, change-scores (Δ = Stim-Pre) were larger in the opsin group compared to the controls (n = 3) for continuous illumination only, with no effects at 5-20 Hz (Fig. 3c). Consistent with behavior, dopaminergic signals were selectively elevated under the efficacious continuous regimen, whereas 5-20 Hz conditions remained indistinguishable from controls (Supplementary Fig. 2d).

**Figure 3:**
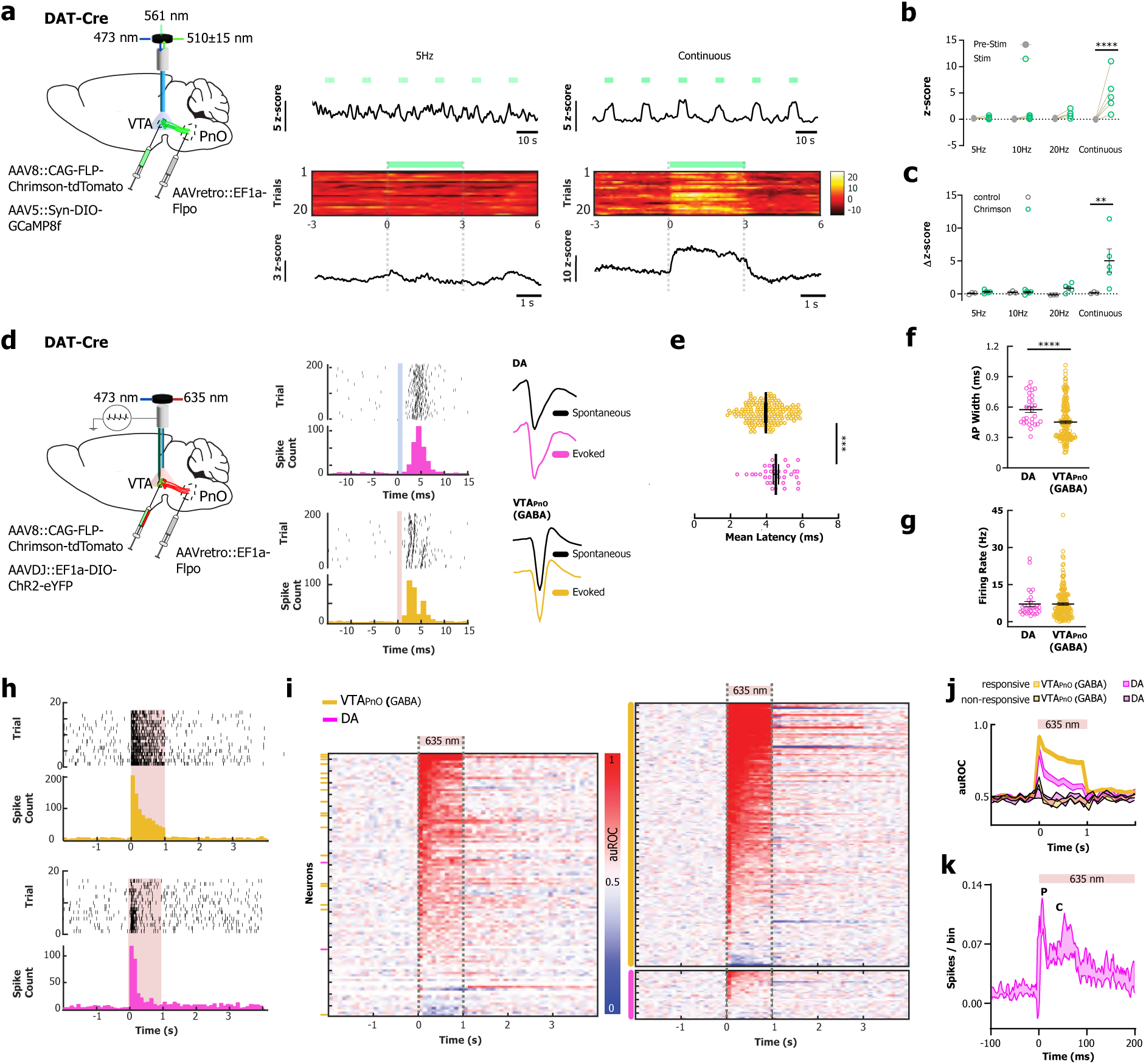
VTA_PnO_ pathway recruitment and DA activity in vivo. **a,** Left, schematic of strategy for fiber photometry with optogenetic activation: DAT-Cre mice received retroAAV-FlpO in PnO and AAV-FLP-Chrimson and AAV-DIO-GCaMP8f in VTA, enabling activation of VTA_PnO_ neurons while imaging local DA population signals. Right, example GCaMP8f traces and peri-stimulus heat maps during 5-Hz trains (3 s) and continuous illumination (green bars). **b,** Baseline (*−*1 to *−*0.1 s) to stimulation (0–3 s) DA population signal comparisons for different train frequencies: continuous (*p <* 0.0001) and 5–20 Hz stimulations (*p >* 0.05, two-way ANOVA with FDR-corrected post hoc comparisons). **c,** DA signal change score (Δ = Stim *−* Pre) comparison between opsin (*n* = 5) and control (*n* = 3) mice: continuous (*p* = 0.002) and 5–20 Hz stimulations (*p >* 0.05, two-way ANOVA with FDR-corrected post hoc comparisons). **d,** Left, schematic of acute opto-tagging and recording strategy: DAT-Cre mice received retroAAV-FlpO in PnO and AAV-FLP-Chrimson and AAV-DIO-ChR2 in VTA with a 64-channel silicon optrode. Right, exemplary opto-tagged neurons: raster/PSTH for a DA unit (magenta) and a VTA_PnO_ unit (GABA, yellow) with waveforms. Light stimulation occurred at time 0 s (1-ms light pulses; colored areas). **e–g,** Summary features of opto-tagged units. Onset latency: DA (4.99 *±* 0.24 ms) vs. VTA_PnO_ (3.82 *±* 0.07 ms) (*p <* 0.0001). Spike width: DA (574.1 *±* 27.3 *µ*s) vs. VTA_PnO_ (451.5 *±* 14.3 *µ*s) (*p* = 0.0002). Firing rate: DA (7.142 *±* 1.02 Hz) vs. VTA_PnO_ (7.162 *±* 0.48 Hz) (*p* = 0.649). Mann–Whitney tests. **h,** Representative opto-tagged responses during continuous 635-nm light for VTA_PnO_ (yellow) and responsive 635-nm light DA (magenta) units. **i,** Left, single-session auROC heat map of all simultaneously recorded VTA units across the silicon probe span (*∼* 1200 *µ*m); side bars mark opto-tagged units (yellow = VTA_PnO_, magenta = DA). Many units show sharp onset modulation, indicating recruitment beyond the opto-tagged subset. Right, pooled auROC heat map of all opto-tagged units across animals (VTA_PnO_ *n* = 173; DA *n* = 30), sorted by onset. **j,** Population responses show reliable, light-locked VTA_PnO_ (GABA) activation and heterogeneous DA outcomes. Responsive DA units exhibited an onset-locked signal transient (*p* = 0.0013, one-sample *t*-test against chance). **k,** PSTH for responsive DA units (2-ms bins). P (peak latency, 7 ms) and C (centroid latency, 41.6 ms) computed over 0–100 ms. Responsive DA units exhibited a consistent early-window activity increase (0–100 ms; baseline-subtracted 0.50 spikes*·*s*^−^*^1^, 95% CI 0.30–0.66; permutation test, *p* = 0.029).

### Acute opto-tagging reveals transient DA activation by VTA_PnO_ optogenetic stimulation

To interrogate projection-defined VTA_PnO_ neurons and local dopaminergic cells at single-cell resolution, we combined cell-type-specific optogenetics with acute high-density silicon probe recordings in DAT-Cre mice (n = 7). To this end, we used a dual viral approach (Fig. 3d left) to target two VTA populations: DAT-positive neurons expressing ChR2 and PnO-projecting neurons (putative GABAergic) expressing Chrimson, which share a dual projection architecture. Following expression, isoflurane-anesthetized mice were recorded with 64-channel silicon optrodes coupled to VTA light delivery, enabling optogenetic identification and simultaneous sampling of both populations. Across seven mice, we isolated 173 opto-tagged VTA_PnO_ neurons and 30 opto-tagged DA neurons (see Methods for opto-tagging classification criteria). VTA_PnO_ units exhibited short-latency, low-jitter light responses; DA units showed longer, more variable latencies and broader spikes, consistent with cell-type identity (Fig. 3e-g; Supplementary Fig. 3b,c). Population analyses confirmed reliable, light-locked firing in optotagged VTA_PnO_ cells. Inspection of a representative recording including all simultaneously sampled VTA units revealed sharp modulation at light onset with differential patterns across the stimulation bout (Fig. 3h) indicating that recruiting the VTA_PnO_ subpopulation propagates activity across the ∼900-1200 *µ*m dorsoventral extent sampled by the probe, beyond the directly tagged units (Supplementary Fig. 3d). Using our activation criterion (per-unit auROC in six 0.5-s windows vs. time-matched baseline; see Methods), a subset of recorded DA units was classified as responsive to VTA_PnO_ stimulation, while the remaining DA units did not meet the criterion (Fig. 3i). Responsive DA units exhibited an onset-locked transient followed by a brief decay; non-responsive units remained near baseline. Within the responsive set, time-binned auROC responses exceeded chance level (Fig 3j). At 2-ms bin resolution, within the set of responsive DA neurons, firing rose within ∼3 ms, peaked at 7.0 ms (IQR 2.5-22.5 ms), and had a centroid of 41.6 ms (IQR 35.5-43.6 ms) with a consistent early-window increase (0-100 ms, baseline-subtracted 0.50 spikes·s-1; Fig. 3k). These features are consistent with transient disinhibition/synchronization mechanisms engaged by VTA_PnO_ recruitment in vivo, including a polysynaptic component (see Discussion).

### Chronic single-unit recordings show direction-selective engagement during reverse locomotion

To test engagement of projection-defined VTA_PnO_ neurons during backward locomotion, we combined the dual-opsin strategy with chronic 32-channel silicon probes and an implanted fiber while mice stood on a rotarod that rotated backward for 2 s at pseudorandom times (n = 3; Fig. 4a, 20 trials; inter-trial 3-15 s). We opto-identified 40 VTA_PnO_ GABA units (Supplementary Fig. 4a). In a representative session that included all simultaneously recorded VTA units we observed a sharp modulation at rotation onset with various patterns thereafter. The ∼900 *µ*m sensing area of the electric probe also covered neurons outside the opto-tagged subset (Supplementary Fig. 4b). Activation was defined using the same criterion as in the acute experiment (see Methods), and temporal structure was quantified in six contiguous 0.5-s windows spanning the rotation and 1 s after offset (W1-W6, 0-3 s) (Fig. 4b). Among opto-identified VTA_PnO_ neurons, 65% (26/40) were classified as activated and 35% (14/40) as non-modulated. Using the W1-W6 scheme, activated units segregated into early-onset 50% (13/26), late-onset 23% (6/26), offset-only 27% (7/26) classes, with class averages tracking rotation timing (Fig. 4c; waveforms in Supplementary Fig. 4b). In a matched subset of bidirectional sessions (one per mouse), population PSTHs (100-ms bins) restricted to reverse-activated units showed sustained elevation during reverse relative to forward across the rotation epoch (Fig. 4d), indicating direction selectivity within the activated pool. As a population summary, considering all opto-tagged VTA_PnO_ units from these sessions (activated and non-modulated), mean response magnitude over 0-3 s (average across W1-W6) was higher in reverse than forward condition (Fig. 4e). A paired-unit analysis revealed a greater fraction of activated bins during reverse and distinct temporal motifs that reorganized across directions; notably, one unit non-modulated in reverse was activated during forward (Fig. 4f).

**Figure 4:**
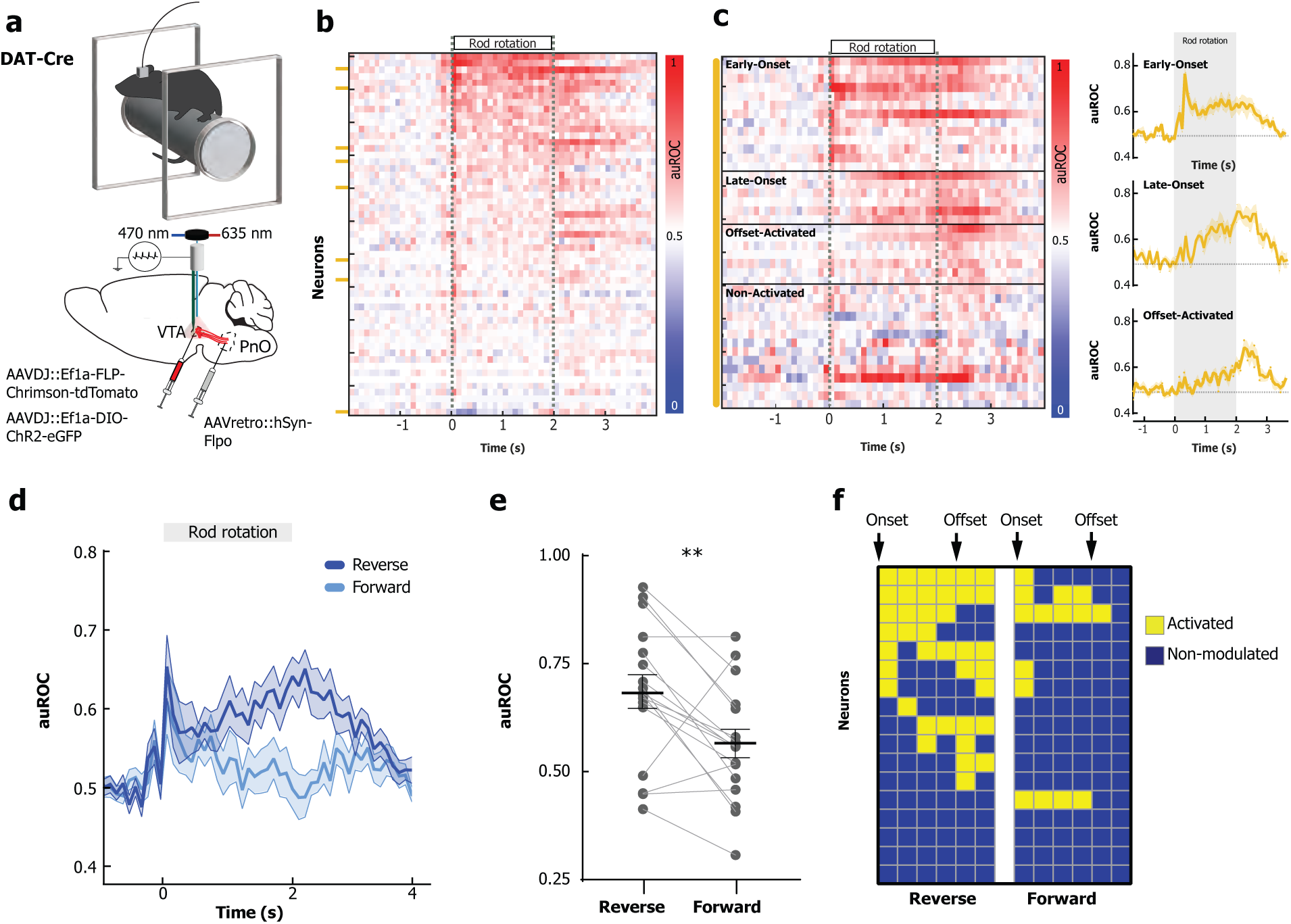
Chronic recordings during forced backward locomotion. **a,** Schematic of chronic recordings strategy: DAT-Cre mice received retroAAV-FlpO in PnO and AAV-FLP-Chrimson and AAV-DIO-ChR2 in VTA with a 32-channel silicon probe and were forced to walk backwards on a rotarod. **b,** Single-session auROC heat map of all simultaneously recorded VTA units across the probe (*∼* 900 *µ*m); side bars mark optotagged VTA_PnO_ neurons (yellow). **c,** VTA_PnO_ activation profiles during rod rotation (0–2 s). Left, heat maps of classified units, sorted by response type: early-onset, late-onset, offset-activated, non-activated. Right, averaged PSTHs for different response classes. **d,** Reverse vs. forward response comparison of reverse-activated VTA_PnO_units (criterion: Methods): population PSTH (100-ms bins) is higher in reverse condition across the rotation epoch (0–2 s), indicating direction selectivity within the responsive pool. **e,** Mean response magnitude of all opto-tagged VTA_PnO_ units over 0–3 s (0–2 s rod rotation and 1 s post stimulus) is higher in reverse than forward condition (paired *t*-test, *p* = 0.0038), summarizing stronger reverse engagement across the population. **f,** Backward vs. forward activation profiles of VTA_PnO_ identified units during forced locomotion.

## Discussion

VTA GABA neurons are often conceptualized as local interneurons that regulate dopaminergic output [5–8]. Recent work, however, demonstrates that long-projection neurons also form intra-VTA synapses, reframing “local inhibition” as a property that can also arise from projection-defined cells, rather than being restricted to a separate interneuron class [13]. Our findings extend this framework by identifying a projection-defined VTA_PnO_ subpopulation that combines long-range output to a brainstem premotor node with monosynaptic local GABA_A_ inhibition within VTA. Functionally, recruitment of this pathway produces coordinated local and distal effects that translate into directionally specific motor output. Thus, rather than operating as purely local modulators or purely long-range effectors, these inhibitory neurons instantiate a dual local–projection architecture that couples midbrain state selection to brainstem motor implementation.

In invertebrates, backward walking is hard-wired by identified descending pathways, and sensory channels can recruit this circuit [28,29]. In mammals, backward stepping appears under broad perturbations or genetic lesions, typically as part of global state changes rather than a selective, instructed program [30–33]. Classical pharmacology experiments showed that concurrent dopamine/serotonin elevation elicits backward walking and circling, whereas receptor blockade suppresses them [19,20]. Thus, backward walking resides in the rodent repertoire but has been engaged mainly by diffuse monoaminergic drives or motor-system instability.

In mammals, descending premotor circuits in the pontomedullary reticular formation implement direction and gait selection [34,35], and basal ganglia outputs can access these brainstem nodes to influence locomotor commands [36]. Within this framework, the ventral tegmental area (VTA) is well positioned to bias direction by engaging premotor effectors; in the present work, we identified a projection-defined VTA inhibitory pathway with both local-projection actions and showed that its activation is sufficient to bias backward locomotion via a defined brainstem target.

Clinically, backward walking is disproportionately impaired in Parkinson’s disease, patients walk more slowly with shorter steps and greater variability, and backward-walking performance shows stronger links to disease severity than forward walking [42,44]. Coordination deficits also emerge during backward walking even when forward-walking metrics are relatively preserved [43]. While we do not claim a disease model, our results motivate testing whether dysfunction in homologous VTA_PnO_ motifs contributes to the selective fragility of backward walking in PD, and whether circuit-level modulation could improve backward walking endpoints used to assess fall risk and progression [42–44].

Our results identify a projection-defined VTA GABA subpopulation with a dual local–projection identity whose activation is sufficient to drive backward locomotion. Optogenetic activation at the soma robustly induced backward locomotion, and selective activation of their terminals in PnO reproduced the effect, supporting a predominantly GABAergic mechanism at a defined brainstem node within the reticulospinal axis [16–18]. Somatic stimulation in VTA produced a faster rise in raw speed than distal activation of VTA_PnO_ terminals (peak *≈* 0.6 s vs. *≈* 1.16 s post-onset), even though both manipulations reached maximal backward-movement proportion at *∼* 400 ms (see Supplementary Fig. 2c). A parsimonious explanation is that driving VTA cell bodies co-recruits local VTA collaterals together with the same long-range axons projecting to PnO, accelerating the motor-state transition; by contrast, selective PnO terminal stimulation engages downstream integration with minimal intra-VTA collateral recruitment and therefore ramps more slowly. This timing dissociation aligns with reports that septal input to VTA scales locomotor vigor/speed and initiates exploration [39–41], and with brainstem premotor nodes that implement speed control [34,35]. The same cells provide monosynaptic GABA*_A_* inhibition locally in VTA, and pathway recruitment elevated population activity with a transient single-neuron increase in a subset of cells, consistent with brief intra-VTA disinhibition coupled to sustained long-range drive. During forced locomotion, tagged VTA-to-PnO units showed stronger engagement during reverse than forward rotations at the population level and displayed distinct, temporally structured direction-selective profiles, distinguishing this pathway from backward walking elicited by diffuse monoaminergic perturbations or ataxic/vestibular instability [19,20,30–33]. Conceptually, these data extend the view that many TH*^−^* VTA neurons combine local collaterals with long-range axons [13]: here, the same inhibitory neurons may coordinate local modulation of DA and non-DA output with distal long-range influence over a premotor brainstem target, providing a circuit mechanism by which basal ganglia outputs access reticulospinal controllers [36] and engage pontomedullary premotor circuits that implement direction and gait selection [34,35].

Although recruiting VTA-GABA neurons is classically expected to suppress DA output [5–8,22], in acute anesthetized recordings optogenetic activation of the VTA to PnO pathway produced a rapid increase in DA single-unit firing (onset ∼3 ms, peak ∼7 ms, centroid ∼38 ms). Two non-exclusive circuit motifs can account for these timescales: (I) Optogenetic phase-locking of the otherwise spontaneous tonic GABA_A_ transients can advance DA spikes via brief hyperpolarization that lowers intracellular Ca^2+^, reduces SK conductance, and permits rebound during the inhibitory trough, without a net reduction in inhibitory tone, yielding sub-10 ms facilitation when inhibition coincides with intrinsic pacemaking [37]. (II) Polysynaptic disinhibition within VTA (GABA-to-GABA) can generate a later boost in DA firing, consistent with the ∼10-50 ms component reflected by the centroid [6]. In this framework, it is plausible that the earliest (3-7 ms) rise is due to synchronous inhibitory gating, with disinhibition contributing to the slower portion of the response.

Differences between the single-unit and photometry readouts likely arise from both sampling geometry and brain state. Our optotagged units were recorded along a narrow mediolateral span but across a deep dorsoventral column (∼900-1200 *µ*m), whereas fiber photometry with a 400-*µ*m, NA 0.55 fiber integrates over a substantially larger near-facet volume; core diameter sets the effective collection field and NA chiefly scales collected power, with most signal arising within ∼200 *µ*m of the facet [45]. Photometry signals are also enriched for neuropil/axonal calcium rather than strictly somatic spikes [46], which can yield sustained fluorescence despite brief somatic spiking. Finally, network dynamics differ across brain states: awake versus anesthetized conditions are known to modulate VTA firing patterns and DA-system dynamics [47]. Together with the kinetics of jGCaMP8f [48], these factors parsimoniously explain the fast-but-sustained DA photometry response versus the brief (∼0-50 ms) DA single-unit activation.

Considering alternative mechanisms, the effect could in principle reflect glutamatergic drive. Although some VTA projection neurons co-release glutamate and GABA [49], our intracellular patch-clamp recordings show that VTA_PnO_ cells make monosynaptic GABA_A_ synapses locally in VTA (IPSCs abolished by picrotoxin/GABA_A_ antagonism), anchoring a GABAergic identity at the soma/collateral level. Thus, while mixed transmitter phenotypes cannot be excluded, the local physiology together with the terminal-stimulation results support a predominantly GABAergic mechanism; the most parsimonious mechanism is local GABA_A_ circuitry plus the timing effects described above [12,38]. Gad2-Cre can label excitatory phenotypes in some regions and is therefore not universally restricted to GABAergic neurons; nonetheless, PnO terminal activation reproduced the behavior and we observed local GABA_A_ IPSCs in VTA, arguing for a GABAergic transmission. Moreover, VTA VGLUT2 maps emphasize ascending outputs (nucleus accumbens, medial prefrontal cortex, lateral hypothalamus, dentate gyrus) and, to our knowledge, do not indicate a prominent PnO projection [11,12,38], making a glutamatergic mechanism at VTA_PnO_ terminals less likely.

The VTA_PnO_ motif described here, with its local-and-projection synaptic architecture, provides a direct link between a midbrain integrator circuit and reticulospinal controllers, offering a tractable entry point to the study of locomotor direction selection.

## Methods

### Juxtacellular recordings/labelings

Adult mice (25-30 g) were anesthetized with urethane (1.25 g kg-1, i.p.). Juxtacellular recordings were obtained from VTA neurons using a dual-channel intracellular amplifier (NeuroData IR-283; Cygnus) and glass micropipettes (tip <1 *µ*m; 6-15 MΩ) filled with (in mM): 250 K-gluconate, 5 KCl, 1 MgCl2„ 2 EGTA, 5 HEPES, 2 MgATP, plus tracer (tetramethyl-rhodamine-biocytin, 2% w/v, or Neurobiotin, 1.7%); pH 7.2. Signals were digitized with a micro1401 A/D converter and acquired in Spike2 v7 (Cambridge Electronic Design). For each neuron we recorded a 200-s baseline (no stimulation), followed by an ipsilateral hindlimb foot-pinch series (3 trials, 15 s pinch per trial; inter-trial interval *≥*60 s). Pinch was delivered with blunt forceps to the plantar surface. After recording, single cells were labeled juxtacellularly according to the standard protocol [21]. Animals were then transcardially perfused with 0.01 M PBS followed by 4% paraformaldehyde in PBS. Brains were post-fixed for 12 h in 4% paraformaldehyde and cryoprotected in 30% sucrose (until sunk), then sectioned at 40 *µ*m on a freezing microtome. For visualization of Neurobiotin-filled neurons, free-floating sections were incubated for 4 h at room temperature in streptavidin-Cy3 diluted in PBS (1:1000; PBS containing 0.5% Triton X-100. After incubation, sections were washed 3 times in PBS, mounted onto glass slides, coverslipped with fluorescence mounting medium, and inspected under a fluorescence microscope to identify labeled neurons.

### Immunofluorescence

Free-floating coronal sections were rinsed 3x in PBS (5 min each). For antigen retrieval (used only for VGAT and VGluT2), sections were incubated in 0.1 M sodium citrate buffer (pH 6.0) at 80 °C for 30 min, then rinsed 3x in PBS. Tissue was blocked/permeabilized for 1 h at room temperature in PBS containing 3% normal horse serum and 0.3% Triton X-100, and then incubated overnight at 4 °C with primary antibodies diluted in the same blocking buffer. After 3x PBS washes, sections were incubated with species-appropriate fluorescent secondary antibodies (Alexa Fluor 488, or 635; 1:1000; Jackson ImmunoResearch) for 4 h at room temperature protected from light, rinsed 3x in PBS, mounted on glass slides, and coverslipped with fluorescence mounting medium. Confocal images were acquired with identical laser and detector settings across conditions. Primary antibodies: Guinea pig anti-Tyrosine Hydroxylase (TH) (1:5000; Synaptic Systems); rabbit anti-VGluT2 (1:2000; Synaptic Systems); rabbit anti-VGAT (1:5000; Synaptic Systems).

### Three-dimensional reconstructions

Neurons selected for reconstruction had fully filled dendritic domains with all dendrites extending to natural tapering ends. Axons, including local collaterals, were traced from their origin at the soma or proximal dendrites, and those leaving the VTA were followed until they merged into the nigrostriatal or medial forebrain bundle pathways, where fluorescence signal typically diminished. Three-dimensional reconstructions were generated from confocal z-stack images using Neurolucida 2017 (MBF Bioscience, USA), with fragments from each section aligned via the Serial Section Manager. Shrinkage correction factors for the somatodendritic domain along the z-axis were applied to account for dehydration and histological processing [50]. Morphometric parameters were obtained using Neurolucida Explorer 2017.

### Axonal projection quantification

Axonal labeling from VTA GABAergic neurons was quantified using an atlas-based workflow. Whole-brain sections were first registered to the Allen Mouse Brain Atlas using ABBA (Allen Brain Atlas Alignment) [51], which enables automated atlas alignment and standardized anatomical annotation across serial sections. Following registration, anatomically defined regions of interest (ROIs) were generated according to atlas boundaries. Fluorescence intensity measurements were then extracted in QuPath v0.5.1 [52] from the registered sections. For each ROI, mean mNeon fluorescence intensity was measured and background-corrected by subtracting the mean intensity of a local background ROI on the same section. Corrected values were averaged per animal and per region, such that each animal contributed a single value per region, thereby avoiding pseudoreplication. Summary values are reported as mean *±* s.e.m. across animals.

### Brain slice preparation

Acute brain slices were obtained following halothane anaesthesia and decapitation. The brain was quickly removed and 250 *µ*m thick horizontal slices containing the midbrain were cut with a VT1200S Leica vibratome (Leica Biosystems, Nussloch, Germany) under chilled low-sodium N-methyl-D-glucamine (NMDG)-based artificial CSF (aCSF) containing (in mM): 92 NMDG, 2.5 KCl, 1.25 NaH2PO4, 30 NaHCO3, 20 HEPES, 25 glucose, 2 thiourea, 5 Na-ascorbate, 3 Na-pyruvate, 0.5 CaCl2, and 10 MgSO4, saturated with 95% O2-5% CO2 (pH 7.3), according to the methods reported by Ting et al. [53]. Thus, slices were maintained in NMDG-based aCSF at 33.0 *±* 0.5°C for 15 min before adding increasing volumes of a 2 M NaCl solution in NMDG-based aCSF every 5 min for 40 min, and finally transferred to a HEPES-based aCSF for long-term storage at room temperature, containing (in mM): 92 NaCl, 2.5 KCl, 1.25 NaH2PO4, 30 NaHCO3, 20 HEPES, 25 glucose, 2 thiourea, 5 Na-ascorbate, 3 Na-pyruvate, 2 CaCl2, and 2 MgSO4 (pH 7.3). After 1h of recovery, slices were transferred to the recording chamber and perfused with aCSF containing (in mM): NaCl 126, NaHCO3 24, glucose 10, KCl 2.5, CaCl2 2.4, NaH2PO4 1.2 and MgCl2 1.2 (95% O2-5% CO2, pH 7.3).

### Ex-vivo electrophysiology

A single horizontal midbrain slice was placed in a recording chamber (volume ∼0.6 ml) on the stage of an upright Nikon Eclipse NF1 microscope (Nikon, Tokyo, Japan) and perfused with aCSF (2.5-3.0 ml per min, 32 °C). Whole-cell patch-clamp recordings were performed with thin-wall pipettes (4-6 MΩ) filled with (in mM):125 KCl, 10 HEPES, 10 EGTA, 5.2 CaCl2, 4 ATP-Mg2, 0.3 GTP-Na3, 10 phosphocreatine-Na2 (pH 7.3 with KOH). VTA DA neurons were identified according to their location between the midline and the medial terminal nucleus of the accessory optic tract (MT) and electrophysiological features, including slow or absent spontaneous firing (<5 Hz) in cell-attached configuration and the presence of a slowly activating inward current (Ih) in response to hyperpolarizing voltage steps, after gaining whole-cell access [54]. Whole-cell electrical signals were recorded with a MultiClamp 700B amplifier using the pClamp 10 software, filtered at 3-4 kHz using the amplifier’s in-built low-pass filter, digitized with Digidata 1440A and computer-saved at 10 kHz sampling rate (all from Molecular Devices, San Jose, CA, USA). Light evoked currents were obtained using the CoolLED pE-300ultra fluorescence microscopy illumination system (CoolLED Ltd, Andover, UK), switched by a computer-controlled TTL input through a Digidata 1440A digital output, using time-controlled flash beams at 480 or 560 nm wavelength. At -60 mV holding potential, the light evoked synaptic currents consisted of fast inward deflections, due to the high chloride-based electrode filling solution, whose GABA_A_ receptor-mediated origin was confirmed by their sensitivity to picrotoxin (100 *µ*M) at the end of most experimental sessions. Moreover, 6-cyano-7-nitroquinoxaline-2,3-dione (CNQX, 10 *µ*M) and D-(-)-2-amino-5-phosphonopentanoic acid (D-AP5, 50 *µ*M) were constantly present in the perfusing solution in order to avoid contamination from ionotropic glutamatergic components. Tetrodotoxin (TTX, 0.5 *µ*M) and 4-aminopyridine (4AP, 100 *µ*M) were also added to the medium, in order to avoid action-potential-dependent polysynaptic components and to increase presynaptic release probability, respectively.

### Stereotaxic injections

Adult male mice (12-16 weeks) were anesthetized with isoflurane in room air (3-4% induction; 1.0-1.5% maintenance) and secured in a stereotaxic frame on a 37 °C heating pad. All coordinates are from bregma. For Gad2 experiments, we made single VTA injections (AP -3.25 mm, ML +0.50 mm, DV -4.00 mm) of either AAVDJ-CAG-DIO-mNeon (tracing; WZ Biosciences) or AAVDJ-EF1a-DIO-hChR2(H134R)-EYFP (optogenetics; Viral Vector Facility (VVF), ZNZ), 300-400 nL per site. For terminal stimulation, Gad2 mice were bilaterally implanted with 200 *µ*m optic fibers (NA 0.39; Thorlabs) aimed at PnO (AP -4.60 mm, ML *±*2.0 mm, DV -4.25 mm) with a lateral 15° approach. For ex-vivo electrophysiology, we injected 200 nL of retroAAV-EF1a-iCre (VVF, ZNZ) in the PnO and 300-350 nL of AAVDJ-hSyn1-DIO-hChR2(H134R)-mCherry (VVF, ZNZ) in the VTA. To target the VTA_PnO_ subpopulation, DAT-Cre mice received AAV8-CAG-FLPX-rc[ChrimsonR-tdTomato] (Addgene #130909) in VTA (AP -3.25, ML +0.50, DV -4.00) and retroAAV-EF1a-FlpO (Addgene #55637) in PnO (AP -4.60, ML +0.85, DV -4.50); these coordinates were used for all PnO viral deliveries. For combined fiber photometry and optogenetics, DAT-Cre mice were injected in VTA with a 300-350 nL cocktail of AAV5-Syn-DIO-jGCaMP8f (Addgene #162379), and AAV8-CAG-FLPX-rc[ChrimsonR-tdTomato], and 150-200 nL of retroAAV-EF1a-FlpO (55637) in PnO; a 400 *µ*m optical fiber (NA 0.66; Doric) was implanted above VTA (AP -3.25, ML +0.40, DV -4.00). Opsin-control mice received the same PnO retroAAV along with VTA injections combining AAV8-CAG-FLPX-rc[ChrimsonR-tdTomato] and AAV-EF1a-DIO-HDM4-mcherry (Addgene #50461). For in vivo electrophysiology/optotagging, DAT-Cre mice were injected with retroAAV-EF1a-FlpO in PnO and a VTA cocktail of AAVDJ-EF1a-DIO-hChR2(H134R)-EYFP and AAV8-CAG-FLPX-rc[ChrimsonR-tdTomato]; for chronic recordings, after 4-5 weeks of expression an optrode (32-channel H8b with integrated 200 *µ*m fiber, NA 0.39; Cambridge Neurotech) was implanted in VTA and mounted on a nano-drive (Cambridge Neurotech). Carprofen (5 mg kg-1, s.c.) was administered perioperatively, and mice were monitored daily during recovery.

### Behavioral Assays

#### Open-Field

Mice explored a 50 *×* 50 cm arena (opaque walls, even ambient illumination). During each 5-min session, 635-nm (VTA stimulation) or 470-nm light (presynaptic stimulation at PnO) was delivered in 3-s bouts every 10 s, via the implanted ferrule. Laser light power was set to 5 mW at the tip of the fiber. Behavior was recorded with an overhead 1080p camera; videos were analyzed offline with DeepLabCut v2.3.8 (DLC) [55, 56]. Videos (30 fps) were tracked in DeepLabCut (nose, back, tail base). Tracks were likelihood-filtered (*p ≥* 0.9), gap-filled, and smoothed (6-frame moving average). From “Back” we computed framewise displacement (movement vector); the body axis was “Tailbase-to-Back” (orientation vector). After a speed gate (*≥* 0.2 units*·*frame*^−^*^1^), frames were classified by the angle between vectors: forward (*<* 60*^◦^*), backward (*>* 120*^◦^*), and sideways (60–120*^◦^*), with sideways split into leftward/rightward by the 2-D cross-product sign. For each 635-nm bout, peri-stimulus windows (*−*1 to +4 s) were binned at 200 ms; within each bin we computed the proportion of frames in each category per trial, then averaged within mouse. Per-bin (1-s) open-field comparisons reported in the main text used Holm step-down across bins.

For the frequency screen (20, 30, 40, 50, 60 Hz and continuous), the same three mice were tested across all conditions in the 50 *×* 50 cm arena. Trials consisted of 3-s light bouts every 10 s. DLC tracking and frame classification followed the pipeline above (likelihood *≥* 0.9; 6-frame smoothing; speed gate *≥* 0.2 units*·*frame*^−^*^1^; forward/sideways/backward by body-axis angle). For each bout we computed the backward ratio as the proportion of frames labeled “backward” within 200-ms bins over *−*1 to +4 s (30 fps). To avoid anticipatory effects, the baseline was *−*1.0 to *−*0.1 s; stimulus windows were 0–1 s (initiation) and 1–2 s (early maintenance). Per trial, the backward ratio was averaged within each window; per mouse, trial means were averaged to one value per frequency and window.

#### In vivo fiber photometry

Freely moving DAT-Cre mice were recorded in a 20 x 30 cm arena using a Doric fluorescence minicube (GCaMP paths: 470-nm excitation 460-490 nm, emission 500-540 nm; isosbestic 405-nm path). Excitation powers at the ferrule tip were ∼70 *µ*W (470 nm) and ∼20 *µ*W (405 nm). The two LEDs were sinusoidally modulated and multiplexed at distinct carrier frequencies (470 nm: 208.616 Hz; 405 nm: 333.786 Hz). Fluorescence collected through the implanted 400-*µ*m, NA 0.66 fiber was detected on a silicon photodiode and digitized; channels were demodulated by the Doric system and exported for analysis. To correct motion/bleach, the 405-nm trace was fit to the 470-nm trace using a robust linear regression with iterative outlier rejection, and the fitted 405 was subtracted from 470 to yield the corrected signal (ΔF/F computed with a safeguarded denominator). Corrected traces were downsampled, aligned to stimulus timestamps, and converted to z-scores using the -1.0 to -0.1 s baseline. For concurrent optogenetics, Chrimson was driven through the same ferrule with ∼150 *µ*W at 568 nm (external trigger). The photometry cube uses separate excitation/emission bands and dedicated return paths for GCaMP, so the 568-nm stimulus is routed away from the green emission detector; in addition, the 568-nm channel was not modulated at the 405/470 carrier frequencies, which avoids lock-in cross-talk. Non-opsin controls confirmed negligible 568-nm influence on the GCaMP channel.

### In vivo opto-tagging

#### Acute recordings

After 4-5 weeks of expression, mice expressing ChR2 (DA; 470 nm) and Chrimson (VTA_PnO_ GABA; 635 nm) were anesthetized with isoflurane (3-4% induction; 1.0-1.5% maintenance) and secured in a stereotaxic frame. A small craniotomy was opened over VTA and a 64-channel optrode (H3 and H10 acute probes, Cambridge Neurotech; 1200 *µ*m recording span) was lowered to VTA (first penetration: AP -3.25 mm, ML +0.50 mm, DV -4.90 mm, tip from bregma). Each experiment comprised 3-4 penetrations, shifting the entry site by ∼200 *µ*m (anterior/posterior/medial) while targeting the same depth. Recordings began with a baseline epoch, followed by cell-type identification using 1 ms light pulses at 5Hz (3 seconds ON 10 Seconds OFF; 210 pulses in total): 470 nm/15mW to tag dopaminergic ChR2+ units and 635 nm/5mW to tag VTA_PnO_ GABA Chrimson+ units. After tagging, we delivered continuous 635 nm light (1 s) every 10 s for 3 min to probe pathway recruitment. At the end of the session, mice were transcardially perfused with 4% paraformaldehyde in PBS.

#### Chronic recordings

Mice with chronic VTA implants (32-channel optrode with integrated fiber; H8b and H10b, Cambridge Neurotech) were connected to the headstage and patch cord and placed in a 20 x 30 cm arena for baseline activity and opto-tagging (1-ms pulses; 470 nm for ChR2-DAT units, 635 nm for Chrimson-tagged VTA_PnO_ units). Animals were then transferred to a custom rotarod with side walls adjusted to ∼1 cm clearance per flank (freely-moving but laterally constrained). The rod was driven in 2-s bouts at 2.00 rad·s-1 (5-cm diameter; surface speed 5 cm·s-1), in the reverse direction such that maintaining position required a backward stepping pattern; inter-bout intervals were pseudo-random 4-35 s (per-mouse mean 9-14 s). Forward sessions (same timing and speed) were run in separate blocks for direction controls; forward and reverse trials were not interleaved. Behavioral video and stimulation TTLs were recorded and synchronized to neural data for peri-event analyses.

### Quantification and statistical analysis

Statistical analyses were performed in MATLAB R2023b, Python 3.10 (statsmodels v0.14.4, SciPy v1.15.1, NumPy v2.2.2, and pandas v2.2.3), and GraphPad Prism v10. All datasets were tested for deviations from normal distribution using Shapiro-Wilk test for appropriate use of parametric data analysis (t-test, ANOVA). When required, non-parametric tests were used (Mann-Whitney, Kruskal-Wallis). Significance level was set at $\alpha$ = 0.05 and p-values were adjusted following multiple comparisons using FDR (False Discovery Rate) or Holm’s step-down corrections. Data is presented as mean *±* s.e.m. Significant differences are denoted in figures as follows: *p < 0.05, **p < 0.01, ***p < 0.001, ****p < 0.0001.

#### Juxtacellular pinch (single-unit and population)

Spike rates were aligned to pinch onset (*t* = 0) and binned at 1 s over [*−*20, +20] s. Baseline was *−*10 to 0 s; Stim was 0–15 s. Responsive units exhibited *≥* 2 consecutive FDR-significant bins during Stim (positive = excited; negative = inhibited; both = mixed).

#### Fiber photometry

Trials were aligned to light onset and averaged within animal to one z-score trace per condition. Epochs: Pre = -1.0 to -0.1 s (excluding the last 100 ms) and Stim = 0.0-3.0 s.

#### Acute silicon-probe recordings (opto-tagging and DA early response)

For each unit, we quantified light-evoked spiking in the early post-stimulus window by summing spikes in the first 6 ms after the light pulse. To assess significance, we built a null distribution by circularly shuffling each trial’s 0–100 ms peri-stimulus spike train 10,000 times and, for each shuffle, taking the maximum 6-ms peak within 0–100 ms. Units were classified as light-responsive when the real 0–6 ms count exceeded the 99.9th percentile of this shuffled distribution. As an inclusion criterion, we required a per-trial response fidelity of *≥* 0.1 (*≥* 1 spike in the 0–6 ms window on at least 10% of pulses; [57]). Third, we also applied a waveform-stability criterion: for each unit, spikes within *−*180 to 0 ms (pre-stim) and 0–6 ms (post-stim, relative to a 1-ms light pulse; 210 pulses) were averaged to yield pre/post mean waveforms, and their Pearson correlation was required to be *≥* 0.90. Units failing any of the three criteria (shuffle test, *≥* 0.1 fidelity, or waveform correlation) were not considered opto-tagged.

#### Acute recordings

classification of responsive vs non-responsive units=. PSTHs were computed (0.5 s bins; -1 to +1.5 s) and compared against a pre-stimulus baseline (-1.25 to -0.25 s). For each bin, auROC quantified the direction and magnitude of change relative to baseline. Bin-wise significance was assessed with paired two-tailed tests versus time-matched baseline and FDR correction across bins. Units were labeled responsive (activated) if they showed a significant increase and an auROC > 0.5; otherwise, they were labeled non-responsive. High-resolution timing (2-ms bins) was used for DA response analysis in acute experiments. Activation status was not re-estimated at 2-ms resolution; DA units inherited their responsive/non-responsive labels from the auROC-based criterion above. The 2-ms analysis quantified onset, peak, and centroid latencies within 0-100 ms and the early-window magnitude (0-100 ms integrated, baseline-subtracted) within the responsive set; non-responsive units were summarized separately as near baseline.

#### Chronic silicon-probe recordings (Direction-selectivity analysis)

Peri-rotation spiking was binned at 100 ms and referenced to a pooled baseline (*−*1.25 to *−*0.25 s) to compute auROC (0.5 = no change). The rotation epoch (0–2 s plus 1 s after offset) was partitioned into six 0.5-s windows (W1–W6). For each unit and window, we compared to a time-matched pre-rotation baseline; *p*-values were FDR-corrected across windows (and directions, when applicable). Significant auROC *>* 0.5 was labeled activation; *<* 0.5 suppression; otherwise, non-modulated. Temporal classes were assigned from reverse-rotation data by first significant window/pattern: Early-Onset (W1, with later significance or run *≥* 2), Late-Onset (W2–W3, with later significance), Offset-Activated (none in W1–W4, *≥* 1 in W5–W6), or Non-Activated.

For direction contrasts (reverse vs. forward; one session per mouse; *n* = 17 units, 3 mice), we ran two complementary analyses: (i) a specificity analysis using only reverse-activated units to compare reverse vs. forward PSTHs and window profiles; and (ii) an all-units population summary (activated and non-modulated) comparing the mean response magnitude over 0–3 s (average of W1–W6) within unit by paired *t*-test, with per-window (0.5 s) reverse/forward heat maps shown in Supplementary Fig. S3. Totals: 40 opto-tagged units overall across 3 mice.

#### Electrophysiology acquisition (opto-tagging) and spike sorting

Acute and chronic extra-cellular recordings were acquired with the Open Ephys GUI v0.6.7 (plugin-based GUI; [58]) using Intan RHD headstages (acute: RHD2164; chronic: RHD2132) at 30 kHz sampling per channel. Signals were digitized at the headstage, streamed via the Open Ephys board, and logged through the Open Ephys GUI. Raw extracellular recordings were imported and preprocessed with SpikeInterface v0.101.0 (Python; [59]), spike sorting was performed with Kilosort v4.0 [60], and the resulting clusters were manually curated in Phy v2.0b5 [61].

#### Open-field video (DLC)

DeepLabCut outputs were likelihood-filtered (*p ≥* 0.9), gap-filled (short dropouts), and smoothed (6-frame moving average). Movement vectors (Back) and body-axis vectors (Tailbase-to-Back) were used to classify frames as forward (*<* 60*^◦^*), backward (*>* 120*^◦^*), and left/right sideways (60–120*^◦^*, cross-product sign) after a speed gate of *≥* 0.2 units*·*frame*^−^*^1^. Per bout, peri-stimulus windows (*−*1 to +4 s) were binned at 200 ms; within-bin proportions of frames per category were computed per trial and averaged per mouse. Group effects used a linear mixed-effects model (Proportion *∼* Time_bin *×* MovementType; random intercept: MouseID). Primary within-frequency inference (per window) used paired tests comparing the per-mouse stimulus mean to its own baseline (paired *t*-test; Wilcoxon signed-rank as robustness). *P* -values across frequencies within a window were adjusted by Holm step-down (family-wise *α* = 0.05). To leverage trial counts while respecting mouse-level dependence, we also modeled per-trial Δ (stimulus *−* baseline) with a linear mixed-effects model: fixed effect frequency (20, 30, 40, 50, 60 Hz; 20 Hz as reference) and a random intercept for mouse (trials nested within mouse). Planned contrasts tested each *≥* 30 Hz level vs. 20 Hz with Holm’s step-down correction; an ordered linear trend (20–60 Hz) was assessed by coding frequency as a numeric factor. Continuous stimulation was compared directly to 60 Hz with the same per-trial mixed-effects framework (reference = 60 Hz).

## Declaration of generative AI and AI-assisted technologies in the writing process

The authors used an AI-assisted language tool for copy-editing (grammar, style, and clarity) of human-written text. All scientific content, analyses, and interpretations were created and verified by the authors. Parts of the analysis code were iteratively debugged/refactored with assistance from an AI tool; all code was reviewed, tested, and validated by the authors.

### Acknowledgements

This research was funded in whole or in part by Aligning Science Across Parkinson’s (ASAP-020505) through the Michael J. Fox Foundation for Parkinson’s Research (MJFF). This work was supported by FONDECYT Postdoctoral Grant 3160763 to C.G.-C. For the purpose of open access, the author has applied a CC BY public copyright license to all Author Accepted Manuscripts arising from this submission.

## Data availability

Source data and key resources used in the current study are available as indicated in the Key Resource table posted at the Zenodo repository at: https://doi.org/10.5281/zenodo.18661699

Standardized in vivo extracellular electrophysiology and fiber photometry datasets are available through the DANDI Archive under Dandiset 001955, version 0.260828.0749: https://doi.org/10.48324/dandi.001955/0.260828.0749

All detailed protocols have been uploaded in the public repository Protocols.io under the accession code: https://www.protocols.io/view/vta-pno-article-hrxcb57ix

Protocols.io collection DOI: https://doi.org/10.17504/protocols.io.bp2l6epmrgqe.v1

The code used for the analyses in this study is available on GitHub at: https://github.com/CGonzC/VTAPnO_Locomotion_Analysis

A citable archived version of the code is available on Zenodo at: https://doi.org/10.5281/zenodo.21938449

Additional requests for information should be addressed to the corresponding author C.G-C.

## Supplementary Figures

**Supplementary Figure 1.**
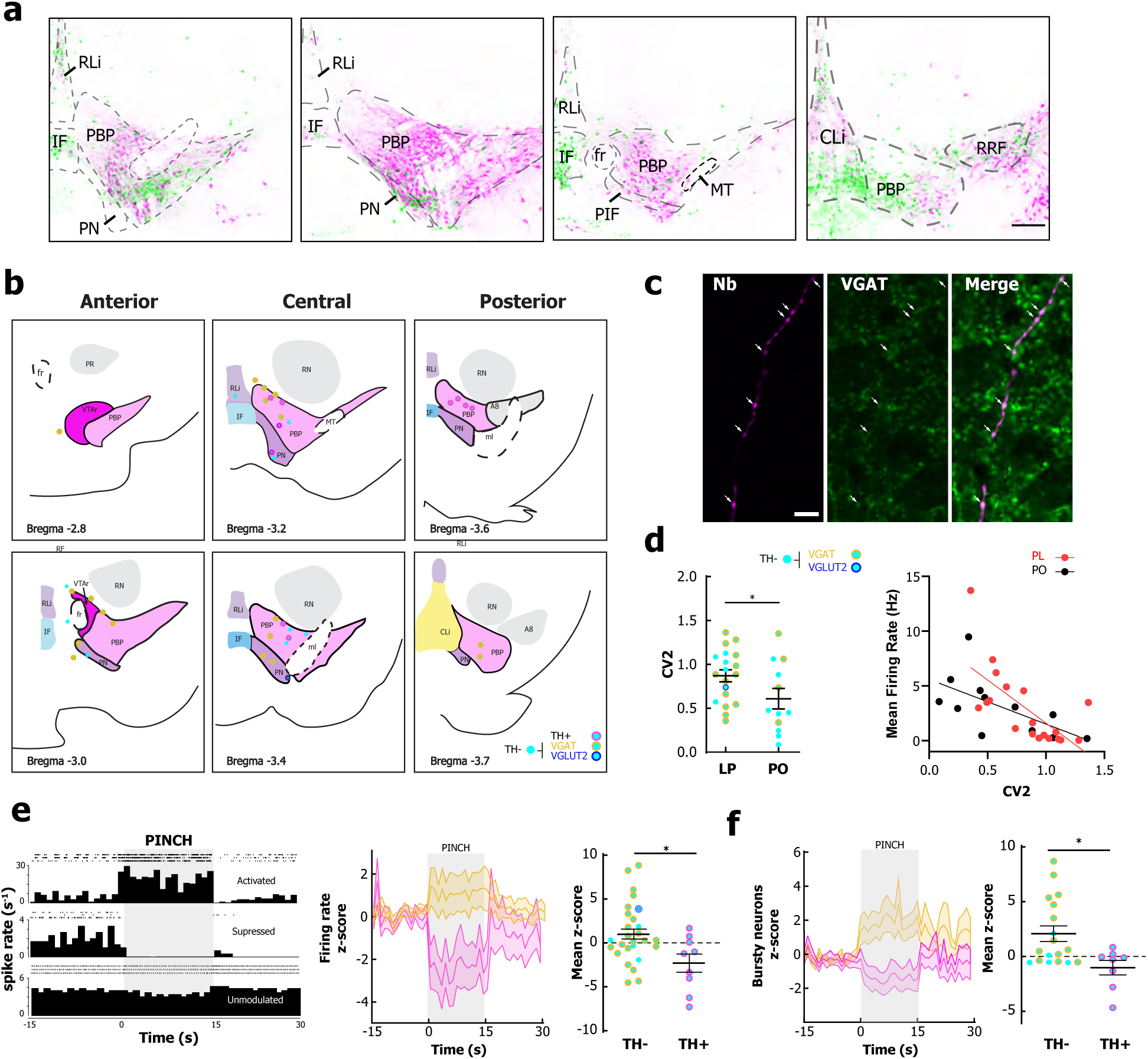
Juxtacellular reconstruction summary and pinch responses. **a,** Example injection spread to visualize GABAergic (in green) VTA cells and their processes across different VTA subfields: RLi, IF, PBP, PN, PIF, CLi, RRF. Magenta: TH immunostaining. **b,** Example reconstructions and anatomical coverage across the VTA (anterior/central/posterior levels; bregma values indicated). **c,** Example VGAT immunostaining in a neurobiotin-filled neuron (Nb). Arrows indicated VGAT-stained boutons. **d,** Left: spike-time variability (CV2) across juxtacellularly labeled TH*^−^* cells: LP (0.871 *±* 0.069) vs. PO (0.609 *±* 0.116) (*p* = 0.0468, unpaired *t*-test). Right: relationship between CV2 and mean firing rate of all recorded TH*^−^* neurons (LP: Spearman *r* = *−*0.76, *p* = 0.0002, *n* = 19; PO: *r* = *−*0.73, *p* = 0.0096, *n* = 12). **e,** Left: hindpaw pinch response profiles (activated/suppressed/unmodulated). Centre: peri-event *z*-scored firing-rate traces of TH*^−^* (yellow) and TH^+^ (magenta) cells. Right: mean firing rate in response to paw pinch is lower for TH^+^ (*n* = 9) than TH*^−^* (*n* = 31) cells (*p* = 0.021, Mann–Whitney test). **f,** Left: pinch-evoked changes in bursting activity of TH*^−^* (yellow) and TH^+^ (magenta) cells (depicted as peri-event *z*-score). Right: mean firing rate of bursting activity in response to paw pinch is lower for TH^+^ (*n* = 8) than TH*^−^* (*n* = 18) cells (*p* = 0.047, Mann–Whitney test). Scale bars: **a,** 200 *µ*m; **c,** 20 *µ*m.

**Supplementary Figure 2.**
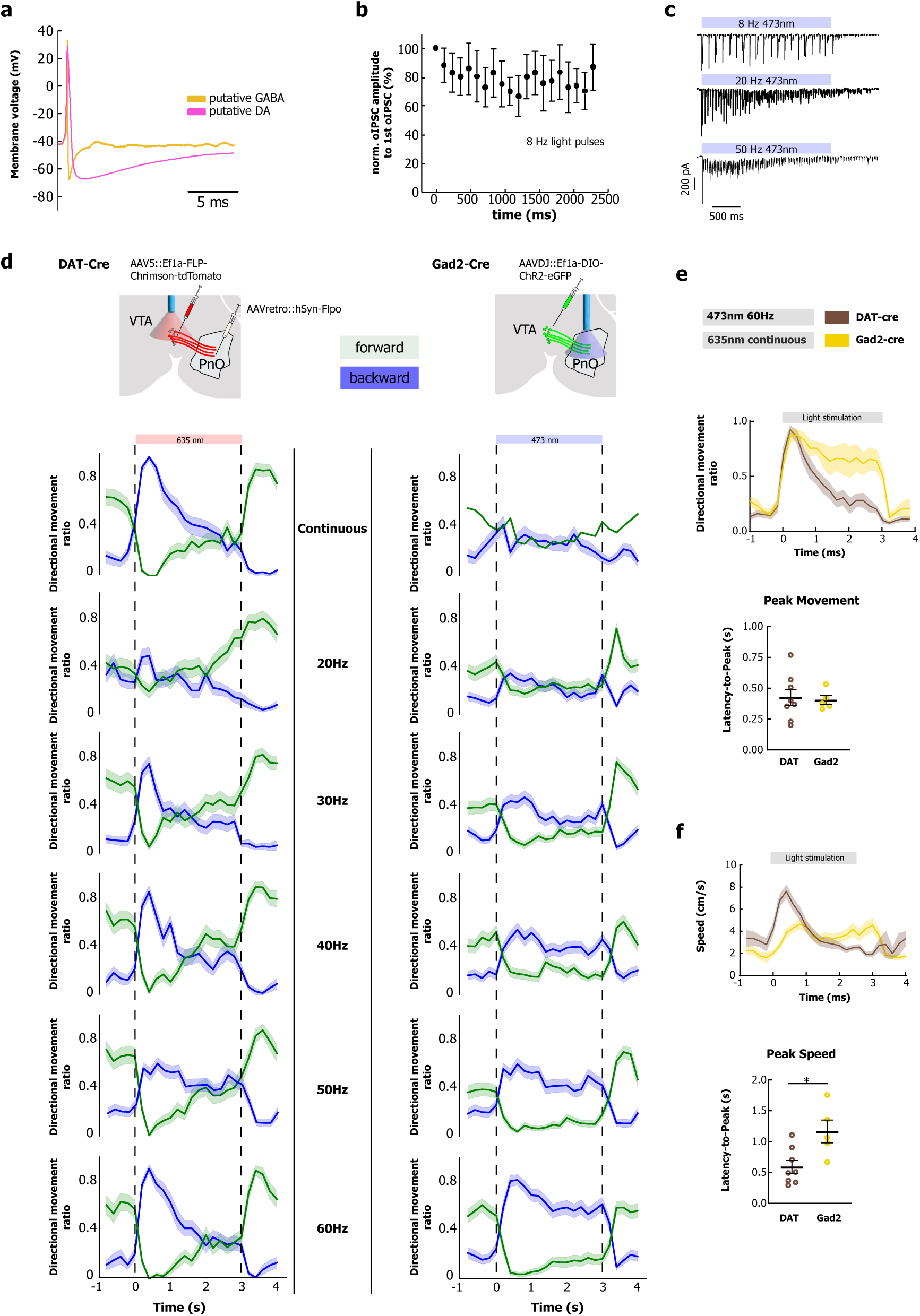
Ex vivo physiology, frequency screen for backward locomotion, and timing metrics. **a,** Example light-evoked responses from putative GABAergic and dopaminergic VTA neurons during VTA-to-PnO pathway stimulation. **b,** Reliability of optically evoked IPSCs across repeated light pulses (normalized oIPSC amplitude over pulse number). **c,** Example traces illustrating light-evoked IPSCs with decreasing amplitudes for 8, 20, and 50 Hz frequencies. **d,** Frequency dependence of open-field directional movement ratio across stimulation conditions (continuous and 20–60 Hz) shown separately for DAT-Cre (somatic VTA_PnO_ stimulation) and Gad2-Cre (VTA_PnO_ terminal stimulation) mice. Mixed-effects model on backward movement ratio showed a significant linear trend across frequencies (*p <* 0.0001). **e,** Upper panel: overlaid peri-stimulus traces comparing DAT-Cre vs. Gad2-Cre directional ratios. VTA_PnO_ somatic stimulation (DAT-Cre) exhibited a sharp onset of backward movement with stimulus onset, followed by a rapid decline. VTA_PnO_ terminal stimulation (Gad2-Cre) caused backward movement across the full stimulation period (0–3 s). Lower panel: both stimulation protocols resulted in a similar latency to peak movement across animals. VTA_PnO_ somatic stimulation (0.425 *±* 0.066 s) vs. PnO terminals (0.404 *±* 0.035 s) (unpaired *t*-test, *p* = 0.8217). **f,** Upper panel: overlaid peri-stimulus traces comparing DAT-Cre vs. Gad2-Cre speeds. VTA_PnO_ somatic stimulation (DAT-Cre) resulted in higher speed during backward locomotion after stimulus onset, followed by a rapid decline. VTA_PnO_ terminal stimulation (Gad2-Cre) resulted in a moderate increase in speed across the full stimulation period (0–3 s). Lower panel: latency-to-peak speed was higher for VTA_PnO_ somatic stimulation (0.590 *±* 0.104 s) than PnO terminals (1.164 *±* 0.184 s) (unpaired *t*-test, *p* = 0.0131).

**Supplementary Figure 3.**
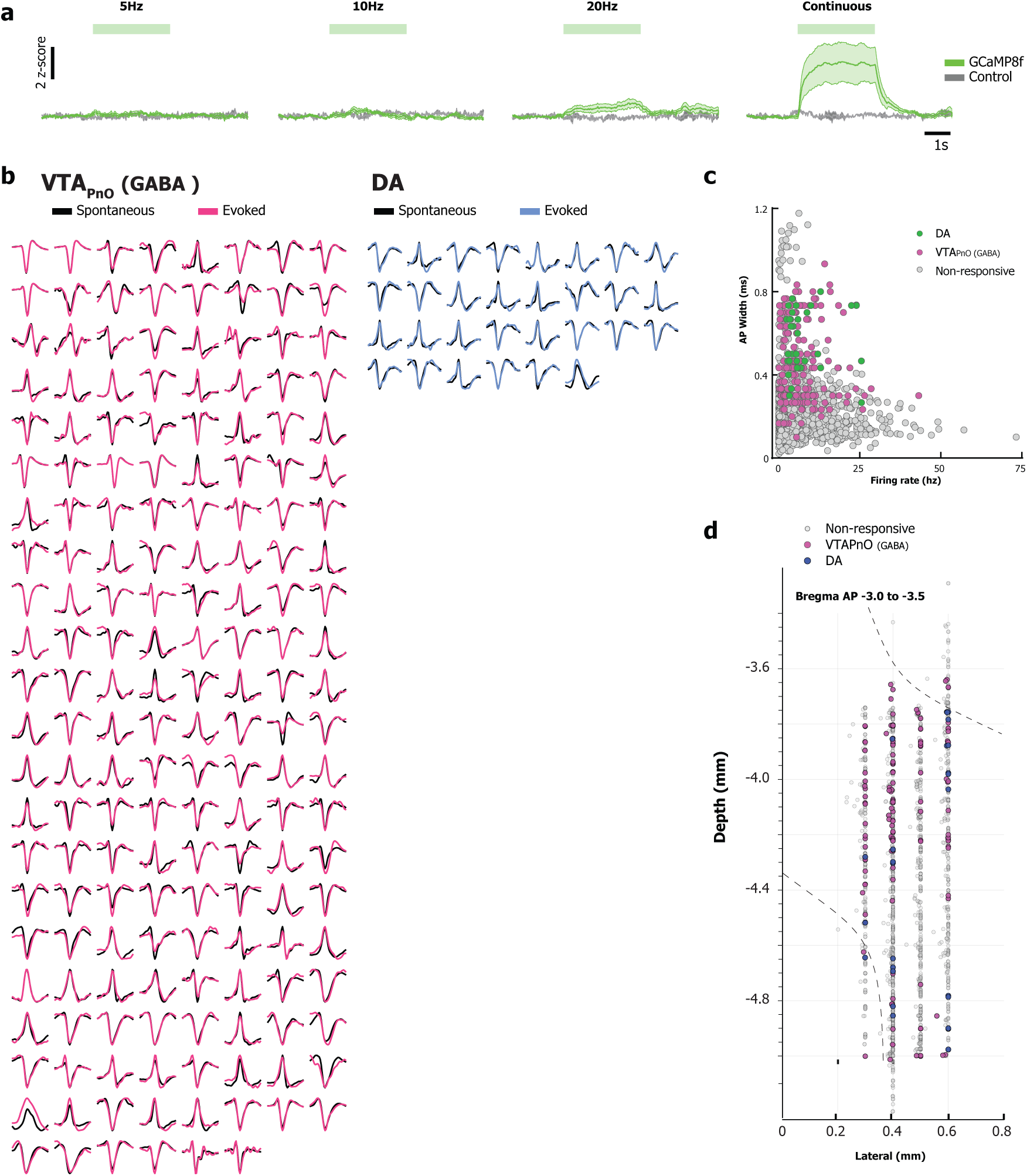
Photometry extended frequencies and opto-tagging feature space/probe coverage. **a,** Fiber photometry (GCaMP8f) peri-stimulus averages across stimulation conditions (5 Hz, 10 Hz, 20 Hz, continuous light), showing opsin vs. control traces (*z*-scored; stimulus delivery indicated by blue horizontal bars). **b,** Waveform examples for opto-tagged units separated by VTA_PnO_ (GABA) and DA, respectively, showing spontaneous vs. optically evoked waveforms. **c,** Feature space scatter plot (firing rate vs. AP width) separating opto-tagged DA units, VTA_PnO_ (GABA), and non-responsive units. **d,** Recording site distributions across silicon probe penetrations (depth vs. lateral position), highlighting responsive categories.

**Supplementary Figure 4.**
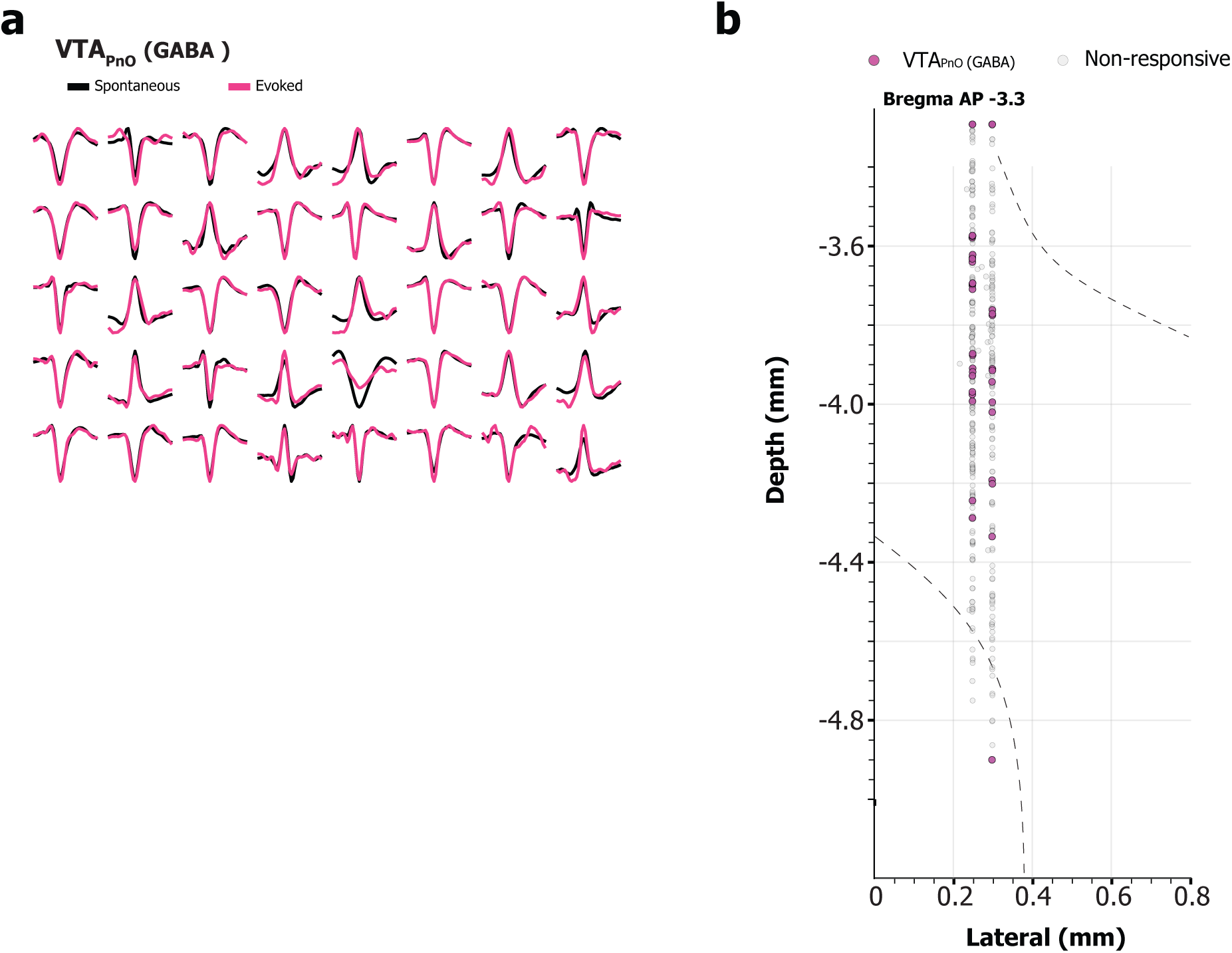
Chronic VTA_PnO_ opto-tagging examples and recording site distribution. **a,** Representative waveforms for opto-tagged VTA_PnO_ GABA units during chronic recordings. Magenta: optically evoked. Black: behavior-evoked. **b,** Chronic recording site distribution (depth vs. lateral position), showing opto-tagged GABA vs. non-responsive units.

